# Mouse thy1-positive spermatogonia suppress the proliferation of spermatogonial stem cells by Extracellular vesicles in vitro

**DOI:** 10.1101/2020.06.15.153668

**Authors:** Yu Lin, Qian Fang, Yue He, Xiaowen Gong, Yinjuan Wang, Ajuan Liang, Guishuan Wang, Shengnan Gong, Ji Wu, Fei Sun

## Abstract

The self-renewal of mammalian spermatogonial stem cells (SSCs) supports spermatogenesis to produce spermatozoa, and this is precisely controlled in a stem niche microenvironment in the seminiferous tubules. Although studies have revealed the role of the surrounding factors in SSCs, little is known about whether the division of SSCs is controlled by extracellular vesicles. Here, extracellular vesicles were found in the basal compartment of seminiferous tubules in mouse, rat, rabbit and human testes. In the mice, the testicular extracellular vesicles are secreted by spermatogonia and are taken up by SSCs. Further, the extracellular vesicles from thy1-positive spermatogonia were purified by anti-Thy1-coupled magnetic beads, and which suppress their proliferation of SSCs but not lead to the apoptosis in vitro.

## 1 INTRODUCTION

Spermatogonial stem cells are undifferentiated spermatogonia that are essential for the maintenance of spermatogenesis. Although SSCs are present in low numbers in the mammalian testis (van den Berg et al., 2007), they balance self-renewal and differentiation to maintain themselves and continually produce committed progenitor spermatogonia, which subsequently become differentiating spermatogonia, spermatocytes, spermatids and mature spermatozoa (Ishii et al., 2012). In human, spermatogonia include progenitor A_dark_-spermatogonia, progenitor A_pale_-spermatogonia, committed A_pale_-spermatogonia, and B-spermatogonia. The progenitor A _dark_-spermatogonia apparently are stems or reserve spermatogonia (Goharbakhsh et al., 2013). In mice, the A_single_, A_paired_ and A_aligned_ spermatogonia are collectively described as undifferentiated type A spermatogonia based on morphological analysis (Clermont and Bustos-Obregon, 1968). These undifferentiated A-spermatogonia undergo a series of cell divisions to form differentiated spermatogonia (Chiarini-Garcia and Russell, 2002a A4, intermediate, and B spermatogonia) before entering meiosis (Chen and Liu, 2015). The A_single_, A_paired_ and A_aligned_ spermatogonia occurs cyclically with in the highly organized seminiferous epithelium, which proliferate primarily in stages I-IV and XI-XII, and remain relatively quiescent at stages of V-X (Sharma and Braun, 2018). Obviously, the SSC division pattern is a complex process and involves positive and negative regulation (Chen and Liu, 2015).

The fate options of SSCs are influenced by extrinsic factors of their stem niche microenvironment comprised of somatic support cell populations that include Sertoli, Leydig, and myoid cells in mammalian testes (Yang et al., 2013). Sertoli cells, the only somatic cell types within seminiferous tubules, physically interact with the SSCs, and likely support SSCs self-renewal by providing growth factors such as glial cell-derived neurotrophic factor (Airaksinen and Saarma) and fibroblast growth factor −2 (Hess et al., 2006; Meng et al., 2000). GDNF supplementation to media is also essential for maintaining the SSCs in culture (Kanatsu-Shinohara et al., 2004). In previous studies, C–X–C motif chemokine −12 secreted by Sertoli cells and colony stimulating factor −1 secreted by Leydig and myoid cells were shown to play critical roles in regulating SSC self-renewal (Chen et al., 2005; Hess et al., 2006; Oatley et al., 2009). Although several positive regulators of SSC self-renewal have been discovered, knowledge of the SSCs niche is still incomplete and much remains unknown about how the microenvironment precisely control the self-renewal division of SSCs, especially the negative regulators.

Extracellular vesicles (EVs) are particles naturally released from the cell that are delimited by a lipid bilayer and do not contain a functional nucleus, EVs are generally referred to as exosomes, microvesicles, and apoptosis body. The term exosome are small (50–100 nm) vesicles that are secreted by a multitude of cell types. Depending on their origin, they can play a variety of roles in different physiological process. It was reported that exosomes are the only class of extracellular vesicles known to be derived from endosomes through the invagination of the endosomal membrane to form multivesicular bodies (MVBs) with numerous small vesicles. These small vesicles are released as exosomes when MVBs fuse with the plasma membrane (Simons and Raposo, 2009). In the mouse testis, Chiarini-Garcia and Russell reported that all type A spermatogonia shokw MVBs, but were not associated with lysosomes (Chiarini-Garcia and Russell, 2002b). As MVB could fuses with the cell surface (the plasma membrane), and release the intraluminal vesicles, which are called exosome, so whether the type A spermatogonia could secrete the extracellular vesicles remained unknown.

Here a large number of EVs (testicular EVs) were found near the basement membrane of seminiferous tubules from rat, rabbit, mouse and human testes across different stages of spermatogenesis, the rat, rabbit, mouse were laboratory rodents that helping us to unravel that a number of EVs in testis appear to be conserved among mammals. We show that the testicular EVs were originated from spermatogonia in the mouse. By separately labelling the testicular EVs, we found that they can be specifically taken up by SSCs, the EVs from thy1-positive spermatogonia were purified by anti-Thy1-coupled magnetic beads, and which suppress their proliferation of SSCs but not lead to apoptosis in vitro. Therefore, we suggest that spermatogonia might use negative feedback to regulate SSC proliferation by secreting EVs.

## 2 MATERIALS AND METHODS

### 2.1 Animals

C57BL/6 mice, rabbit, and adult Sprague Dawley rats were purchased from SLAC Laboratory Animal Co., Ltd. (Shanghai, P. R. China), All animal care and experiments of this study were performed in accordance with the guidelines and were approved by the Ethics Committee of International Peace Maternity and Child Health Hospital, School of Medicine, Shanghai Jiaotong University (Permit number: GKLW 2017-31).

### 2.2 Human testicular biopsies

The collection of human testicular tissues was in accordance with institutional guidelines, and **all patients provided written informed consent**, the study design was approved by the Ethics Committee of the International Peace Maternity and Child Health Hospital. Informed consent was obtained from the participants. Two testicular tissue biopsies were obtained by puncture from men with obstructive azoospermia (age 30 and 37 years) with normal spermatogenesis. Clinical examinations included the evaluation of secondary sexual characteristics, testicular size and consistency, epididymal distension, presence of the vasa deferentia and varicocele. Both patients had their serum follicle stimulating hormone concentrations measured, with values in the normal range.

### 2.3 Cell culture

Spermatogonial stem cells were established from 6-day-old male F1 progeny of DBA/2 × C57BL/6 or C57BL/6/Tg14 (act-EGFP-OsbY01) mice as described (Gong et al., 2017; Kanatsu-Shinohara et al., 2003). There is no difference when it comes to the percentage and behaviour of the SSCs within the testes of C57BL/6 and DBA/2 mouse (Kubota and Brinster, 2008). They were seeded on mitomycin C-treated mouse embryonic fibroblast (MEF) feeder cells and cultured in SSC medium consisting of StemPro-34 SFM medium supplemented with stemPro supplement (Thermo Fisher Scientific, Waltham, MA, USA), 25 μg/ml insulin, 100 μg/ml transferrin, 60 mM putrescine, 30 nM sodium selenite, 6 mg/ml D–(+)-glucose, 30 μg/ml pyruvic acid, 1 μl/ml D-L-lactic acid (Sigma-Aldrich, St Louis, USA), 5 mg/ml bovine serum albumin (Sigma-Aldrich, St Louis, USA), 2 mM L-glutamine, 10 μM 2-mercaptoethanol (Sigma-Aldrich, St Louis, USA), 1 × MEM vitamins solution (Invitrogen, USA), 1 × non-essential amino acid solution (Invitrogen, USA), 2 mM L-glutamine (Invitrogen, USA), 1 × penicillin/streptomycin solution (Invitrogen, USA), 0.1 mM ascorbic acid, 10 μg/ml d-biotin (Sigma-Aldrich, St Louis, USA), 20 ng/ml recombinant human epidermal growth factor (Invitrogen, USA), 10 ng/ml human basic FGF (Invitrogen, USA), 10 ng/ml recombinant human GDNF (Invitrogen, USA) and 1% fetal bovine serum (FBS) (Gibco/Life Technologies, Thermo Fisher Scientific, Waltham, MA, USA). The medium was replaced every 2–3 days. For MEF preparation, C57BL/6J mouse embryos were minced, digested with trypsin-EDTA (Invitrogen, USA), and then cultured in Dulbecco’s (D) MEM containing 10% FBS supplemented with 2 mM glutamine, 100 U/ml penicillin and 100 μg/ml streptomycin (MEF culture medium).

### 2.4 Isolation and analysis of testicular EVs

Twenty mice at 20 days post-partum were euthanized by cervical dislocation, the testes were dissected and decapsulated, then digested with 0.25 mg/ml collagenase IV (Sigma-Aldrich, St Louis, USA) in DMEM/F12 medium at 37 °C for 20 min with slow shaking (150 cycles per min) until the tubules had dispersed fully. The tubules were allowed to settle for 2 min at room temperature by standing the tube vertically. The supernatant enriched in interstitial testicular cells was discarded, leaving just the settled tubules. These were digested with 2 mg/ml collagenase IV and 2 mg/ml hyaluronidase (Sigma-Aldrich, St Louis, USA) at 37 °C for 20 min, centrifugated at 3,000 × g for 30 min to remove the cells, then centrifuged at 10,000 × *g* for 30 min at 4°C to remove cell debris, and the final supernatant was centrifuged at 100,000 × *g* for 90 min in an SWT32 swinging bucket rotor (Beckman Coulter, Brea, CA, USA) at 4 °C to precipitate the EVs. The extracellular vesicle pellets were washed with phosphate-buffered saline (PBS), and filtered using 0.22 µm filter (Millipore, USA), and EVs were finally collected by centrifugation at 100,000 × *g* for 90 min at 4°C and resuspended in 20 μl of PBS. The purified EVs were analyzed using Zeta Particle Metrix equipment (Particle Metrix GmbH, Meerbusch Germany), after which they were stored at −80 °C. The pellets were resuspended in PBS for subsequent analysis.

### 2.5 Purification of Thy1 positive EVs

The isolation of Thy1 positive EVs is performed by positive selection using anti-CD90 MicroBeads (Miltenyi Biotec, USA) according to the manufacturer’s instructions. Briefly, the testicular EVs were diluted with 2 ml phosphate-buffered saline (PBS) containing 0.5% bovine serum albumin, and 2 mM EDTA, and incubated with anti-CD90 beads (100 ul) for 3 hours at 4 °C under slow rotation, A microcolumn (LS separation columns, MACS, Miltenyi Biotec) was placed in MACS magnetic separator and the column was rinsed thrice with 1 mL rinsing solution (MACS BSA Stock Solution diluted 1:20 with autoMACS Rinsing Solution, Miltenyi Biotec). Beads bound to EVs were applied onto a magnetic column and Thy1-negative EVs that passed through the column were collected.

### 2.6 Testicular extracellular vesicle labeling and uptake assay

Purified EVs were labeled with Exo-Glow extracellular vesicle labeling kits (SBI Biosciences, Palo Alto, CA, USA) according to the manufacturer’s instructions. Aliquots of 50 μl of 10 × Exo-red were added to 500 μl of resuspended EVs in PBS, mixed well by flicking the tube and incubated at 37°C for 10 min. The labeling reaction was stopped by adding 100 μl of ExoQuick-TC reagent and incubated on ice for 30 min. The labeled EVs were centrifuged and washed with PBS three times. SSCs were incubated with the labeled EVs for 1–4 h and visualized using fluorescent microscopy (Leica, Wetzlar, Germany)

### 2.7 Cluster-forming activity (CFA) assay

CFA assays were performed as described (Yeh et al., 2007). SSCs were harvested from an established cluster culture and seeded at approximately 1 × 10^4^ cells/cm^2^ in 96-well culture dishes. After incubation with 0, 2, and 4 μl testicular EVs (5 × 10^9^ particles/μl) per well for 7 days, the medium was changed to SSC culture medium. All clusters in a well were counted visually at the 6th day. Experiments were performed in triplicate.

### 2.8 Reverse transcription quantitative polymerase chain reaction (RT–qPCR) amplification

SSCs were separated from MEFs by gentle pipetting and total RNA was extracted from SSCs using TRIzol (Invitrogen, USA) according to the manufacturer’s instructions. Reverse transcription was performed using PrimeScript RT Master Mix (CD201-2, TaKaRa, Otsu, Japan), and qPCR was performed using SYBR Premix Ex Taq II (RR820L, TaKaRa, Japan) in an ABI 7500 Real-Time PCR System (Applied Biosystems, Foster City, CA, USA). The qPCR conditions were 95 °C for 5 min, followed by 40 cycles of 95 °C for 5 s and 60 °C for 34 s. Transcript levels were normalized to the housekeeping gene *Gapdh*.

### 2.9 Western blot analysis

Testicular EVs were lysed in RIPA buffer (P0013B; Beyotime Biotechnology, Shanghai, P. R. China), the protein concentrations were measured using Bradford Protein Assay kits (P0006; Beyotime Biotechnology, China), and exosomal amounts loaded for western blotting were normalized according to the protein concentration. The exosomal lysates were fractionated in 12% sodium dodecyl sulfate polyacrylamide gel electrophoresis (SDS–PAGE), electrotransferred to nitrocellulose membranes blocked with 5% (w/v) nonfat dry milk and incubated with primary antibodies against CD81 (18250-1-AP, Proteintech Group, Wuhan, P. R. China)(diluted as 1:2500), CD9 (ab92726, Abcam, Cambridge, UK) (diluted as 1:2500), Thy1 (ab225, Abcam, Cambridge, UK) (diluted as 1:2500), Gfra1(ab186855, Abcam, Cambridge, UK) (diluted as 1:2500) in 0.1% Tween-20 in tris-buffered saline (TBST) overnight at 4 °C After washing with TBST, the membranes were incubated with secondary antibodies: horseradish peroxidase (HRP)-conjugated goal anti-rabbit IgG (H+L) (SA00001-2, Proteintech, China) (diluted as 1:5000) or HRP-conjugated goat anti-mouse IgG (H+L) (SA00001-1, Proteintech, China) (diluted as 1:5000) in TBST for 1 hour at room temperature. After washing, Immunoblots were visualized using ECL substrate (Thermo Scientific, USA) and ImageQuant LAS4000 Mini software (GE Healthcare Life Sciences, Chicago, IL, USA).

### 2.10 Transmission electron microscopy (TEM)

Testicular samples from rat, mouse, rabbit and human were fixed in 2.5% glutaraldehyde in 0.1 M HEPES buffer (pH = 5) overnight at 4 °C then postfixed in 1% osmium tetroxide. The samples were then rinsed with PBS followed by dehydration in an ethanol gradient and then embedded in Epon 812 (Sigma-Aldrich, St Louis, USA). Ultrathin sections obtained using an Ultracut R ultramicrotome (Leica) were stained with uranyl acetate and lead citrate. For immunoelectron microscopy, mouse testes were fixed in 4% paraformaldehyde/0.2% glutaraldehyde in 100 mM sodium phosphate, pH 7.4, at 4 °C by immersion for 3 h, After washing with 100 mM lysine in 100 mM sodium phosphate, pH 7.4, and 150 mM sodium chloride, they were dehydrated in a graded series of cold ethanol then embedded in Epon 812. Ultrathin sections was incubated with primary antibodies against IgG (30000-0-AP, Proteintech, China), GFRα1 (ab186855, Abcam, Cambridge, UK) for 12 h at 4 °C. Then, 10 nm of colloidal gold-conjugated second antibody (A-31566, Thermo Fisher Scientific, Waltham, MA, USA) was incubated for 2 h at room temperature. After staining with uranyl acetate and lead citrate, images were captured using a transmission electron microscope (H-7560; Hitachi, Tokyo, Japan) at 80 kV.

EVs were fixed in 4% paraformaldehyde and layered on Formvar-carbon-coated electron microscopy grids, washed with PBS, and further fixed with 1% glutaraldehyde for 5 min. Samples were then stained with 4% uranyl acetate for 30 min, after which images of the micrographs were captured using a transmission electron microscope (H-7560; Hitachi, Japan) at 80 kV.

### 2.11 Immunofluorescence Staining

After 5 days of culture, OCT4, PLZF and MVH were used as the target molecule for the identification of SSCs. Briefly, cells were fixed in 4% paraformaldehyde solution (PFA) for 30 minutes, and washed with PBS three times, each time for 5 minutes. Then, cells used for OCT4 and PLZF staining were treated with 0.5% Triton X-100 for 30 minutes at room temperature, cells used for MVH staining should not be treated with Triton X-100. Following washing with PBS for three times, cells were blocked with goat serum at 37°C for 20 minutes, then incubated overnight at 4°C with rabbit-anti-OCT4 (1:100, Santa Cruz Biotechnology), mouse-anti-PLZF (1:100, Santa Cruz Biotechnology) or rabbit-anti-MVH (1:100, Santa Cruz Biotechnology). After that, the cells were washed with PBS containing 0.05% Tween-20, followed by the incubation with Rhodamine (TRITC)-conjugated goat anti-rabbit IgG(H+L) (ProteinTech, USA) or Rhodamine(TRITC)-conjugated goat anti-mouse IgG(H+L) (ProteinTech, USA) for 30 minutes in darkness at 37°C. Next, the nuclear was stained by Hoechst33342. Finally, images were photographed under a DM2500 fluorescence microscope (DMI3000B; Leica).

### 2.12 EdU (5-Ethynyl-2’-deoxyuridine) Assay

For analysis of the effect of EVs on SSCs proliferation, the Cell-LightTM EdU Apollo567 In Vitro Kit (RiboBio, Guangzhou, China) was used according to the manufacturer’s instructions. SSCs were cultured in 96-well plates for 5 days with different treatment (control, Thy1 po EVs and Thy1 ne EVs), followed by the incubation with 50 µM EdU at 37°C for 2 hours, then cells were washed twice with PBS, each time for 5 minutes. After being fixed by 4% PFA for 30 minutes at room temperature, cells were orderly incubated with 50 µL 2 mg/mL glycine solution for 5 minutes and then treated with 0.5% Trion X-100 solution for 10 minutes on a shaker. Subsequently, cells were washed once with PBS and incubated with 1x Apollo staining solution for 30 minutes in darkness on a shaker. After three times of wash with PBS solution containing 0.5% Triton X-100, cells were incubated with Hoechst33342 for 10 minutes at room temperature to stain cell nucleus. Finally, images were obtained under the Leica fluorescence microscope.

### 2.13 Cell Apoptosis Assay

Cell apoptosis detection was performed using Annexin V-FITC Apoptosis Detection Kit (eBioscience, BMS500FI-300) according to the manufacturer’s instructions. After 5 days of culture, SSCs were collected by trypsin digestion and washed once in PBS by gentle shaking or pipetting up and down, SSCs were resuspended in binding buffer. Then, cells were incubated with Annexin V-FITC for 10 minutes at room temperature, after that, SSCs were washed once in binding buffer and resuspended in binding buffer containing 20 µg/mL propidium iodide (PI), and incubated for 15 minutes at room temperature in darkness. Cell apoptosis were analyzed by Beckman Cytoflex (Beckman Coutler Co.Ltd &Cytoflex).

### 2.14 Statistical analysis

Experiments were run in triplicate. Two-tailed, unpaired Student t-test was used for statistical analysis, data are presented as the mean ± standard error of the mean (SEM). *P* < 0.05 was considered to be statistically significant. The statistical graphs were generated by GraphPad Prism 6.

## 3 RESULTS

### 3.1 Testicular EVs are secreted by spermatogonia

To explore the role of EVs in the development of spermatogenesis, we tracked the EVs in the testes from mice at 8 days, 14 days, 21 days, and 8 weeks post-partum (P8, P14, P21, 8w) using TEM. A number of small EVs were found near the basement membrane of seminiferous tubules at all stage, and not in the adluminal compartment, The number of smaller vesicles varies greatly in each cluster, some structures contain up to 150 vesicles, some contain fewer than 20, these testicular EVs had diameters of 30–100 nm (Fig. 1A). In addition, One MVB was detected in the cytoplasm of a type A spermatogonium, (Fig. 1B), and the EVs are released from cells upon fusion of MVB with the plasma membrance, The type A spermatogonia was distinguished on the basis that it contains several compact nucleoli (Fig. 1B arrow); patches of heterochromatin (Fig. 1B arrowhead) are sparse along the nuclear envelope that consistent with previous report(Chiarini-Garcia and Russell, 2002b). Together, these images prompted us to consider whether the testicular EVs were from spermatogonia, to determine that, we separated the testicular EVs by two-step enzyme digestion and differential ultracentrifugation. Western blot analysis of them revealed that they were all detected together with spermatogonial membrane proteins GFRα1, THY1, CD9 and exosomal protein CD81 (Fig. 1C). Further, immunoelectron microscopy showed that GFRα1 protein was detected on testicular EVs in the testis sections from mice at 8w (Fig. 1D). These results indicate that testicular EVs were secreted by spermatogonia.

**Figure. 1.**
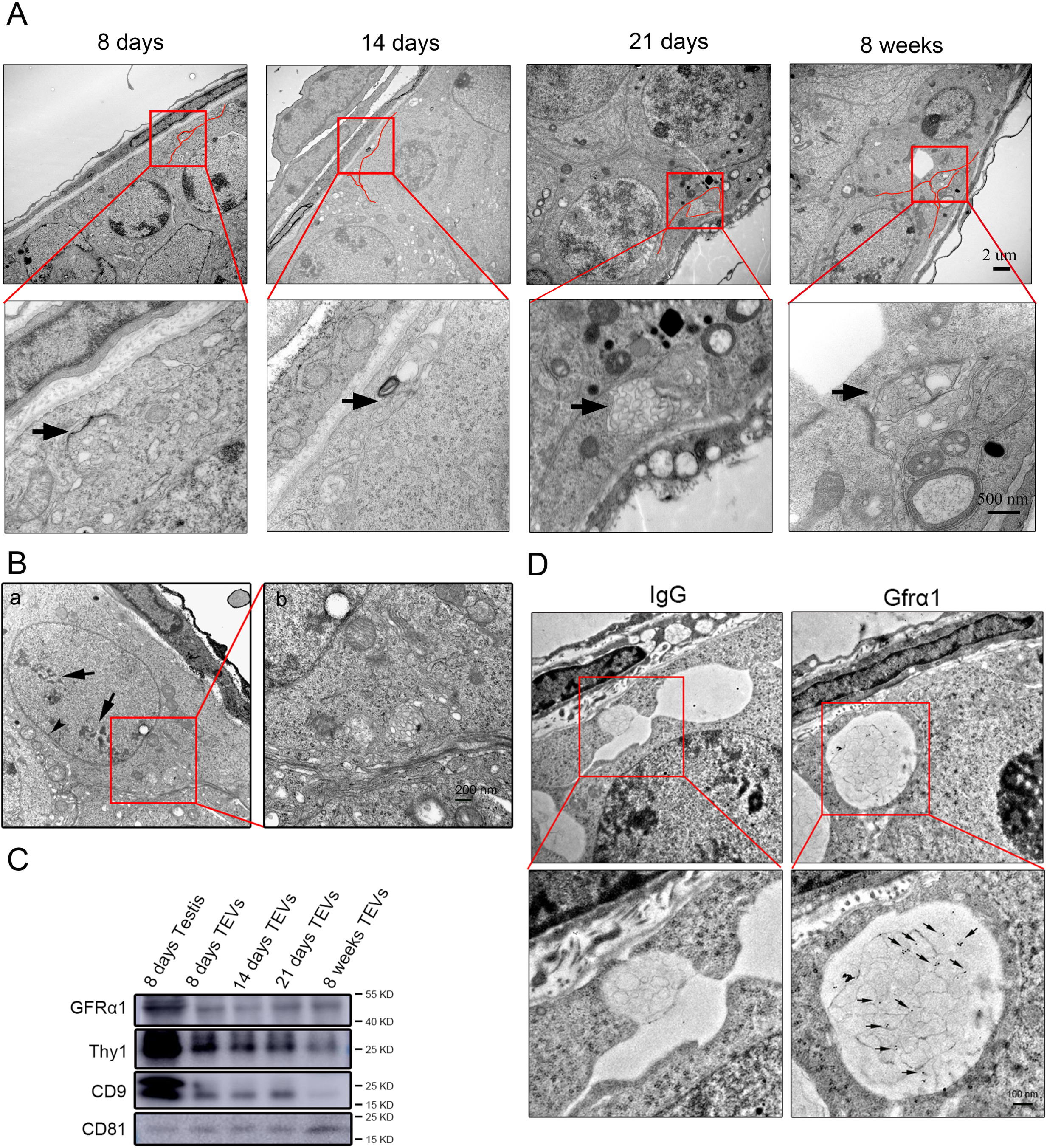
Testicular EVs are secreted by spermatogonia. **A**) TEM images of testicular EVs in cross-sections of seminiferous tubule from mice at 8, 14, and 21 days, and 8 weeks post-partum (P8, P14, P21, and P8w), the number of smaller vesicles varies greatly in each cluster, some structures contain up to 150 vesicles, some contain fewer than 20, the diameter of the testicular EVs ranges from 30 to 100nm, arrow: a number of EVs, red line: plasma membrane. **B**) Transmission electron microscopy (TEM) images of type A spermatogonia from 8-week-old mice, multivesicular bodies (MVBs) were detected in the cytoplasm (**b**). arrow: compact nucleoli, arrowhead: patches of heterochromatin. **C**) Western blot analysis of GFRα1, THY1, CD9, and CD81 expressed in testicular EVs from mice at P8, P14, P20, and 8w. **D**) Immunoelectron microscopy images of testicular EVs from 8-week-old mice showing immunogold labeling for GFRα1 protein was detected in testicular EVs; IgG was used as a negative control, arrow: GFRα1.

### 3.2 EVs are present in mouse, rat, rabbit and human testes

We next asked whether the testicular EVs could be detected in other mammals. To test this, we used TEM to investigate the EVs in rat, mouse, rabbit and human testes. The results shown that testicular EVs were present in the rat, mouse, rabbit and human testes, and they are also closed to the basement membrane of seminiferous tubules and show up in large numbers, these testicular EVs had the same diameters (30–100 nm)(Fig. 2), these results unraveled that a number of EVs in testis appear to be conserved among mammals.

**Figure. 2.**
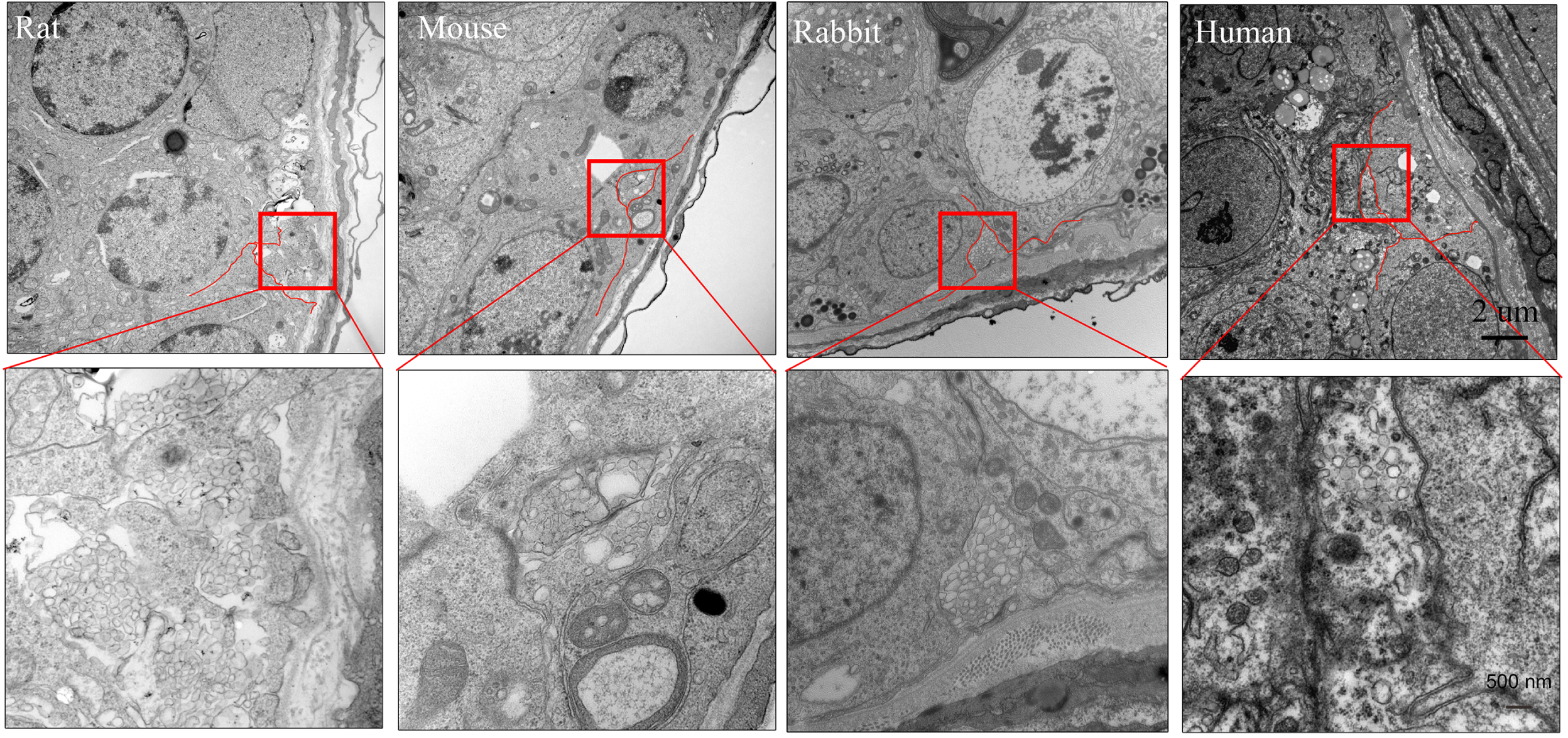
Testicular EVs are present in mouse, rat, rabbit and human testes. TEM images of testicular EVs in cross-sections of seminiferous tubules from rat, mouse, rabbit and human testes, red line: plasma membrane.

### 3.3 Testicular EVs suppress the proliferation of SSCs cultured in Vitro

To explore the function of testicular EVs in spermatogenesis, we first separated the testicular EVs from P20 testes by two-step enzyme digestion, and the second digestive supernatant of seminiferous tubules was used to isolate the EVs by an established ultracentrifugation protocol (Thery et al., 2006). To confirm EVs purification, samples were examined using Nanoparticle tracking analysis (NTA), TEM, and western blotting. Both NTA and TEM of the extracellular vesicle fractions revealed that the diameter of Evs ranged from 50 to 120 nm (Fig. 3A, B). Western blot analysis of extracellular vesicle fractions confirmed the presence of the exosomal proteins CD81 and CD9, as identified in the ExoCarta database(Mathivanan et al., 2012), and the supernatant was not detected the CD81 and CD9 (Fig. 3C).

**Figure 3.**
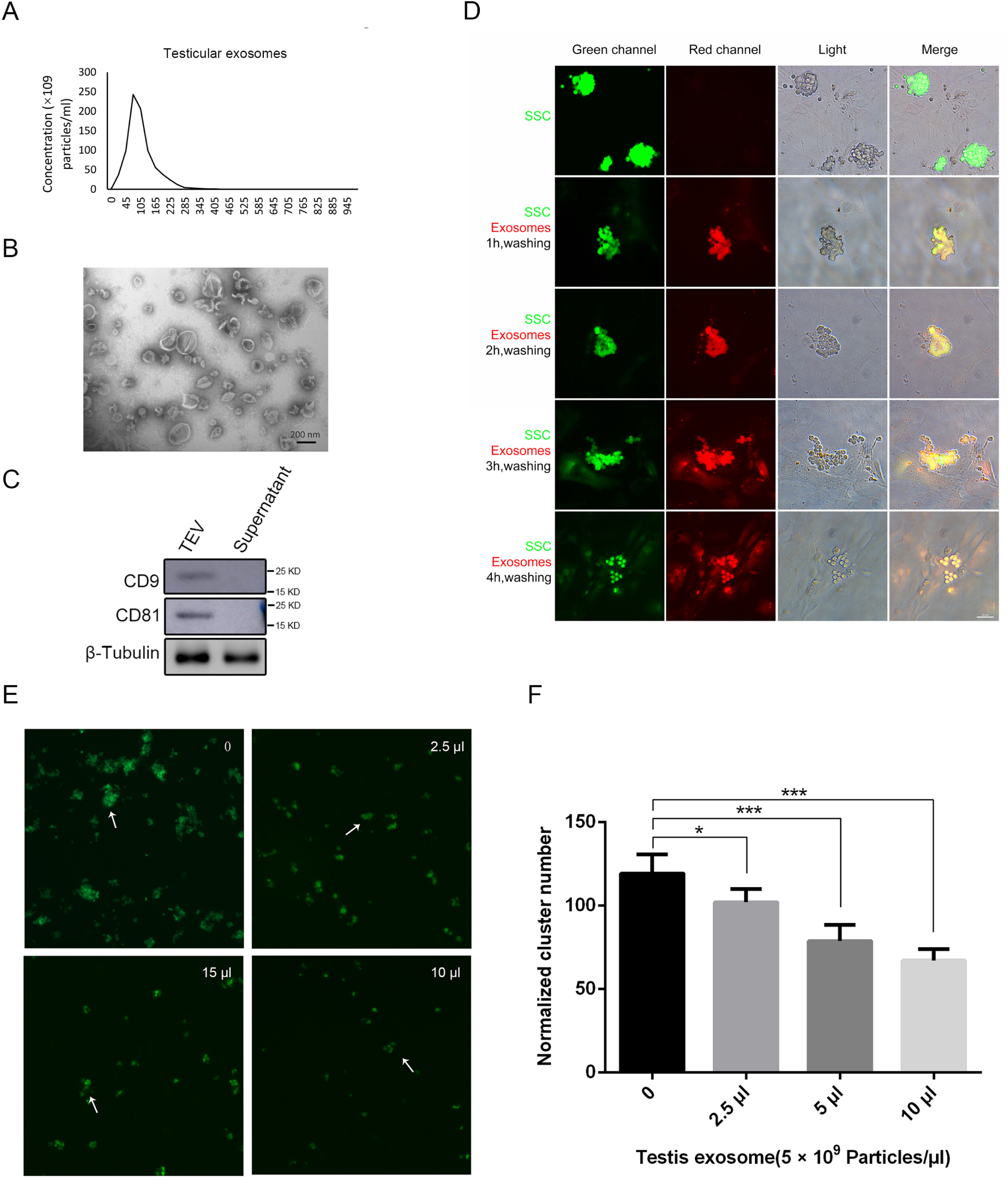
Testicular EVs repress the proliferation of SSCs in vitro. **A**) Size distribution of the testicular EVs determined by NTA analysis. **B**) Representative TEM images of isolated testicular EVs. C) Western blot analysis for CD81, CD9, and beta-Tubulin using extracts from testicular EVs and supernatant. **D**) EVs uptake was visualized using fluorescence microscopy after treatment with Exo-red labeled testicular EVs. **E, F**) The effect of increasing concentrations of testicular EVs on SSCs after 7 days were evaluated from cluster counts. Student’s t tests were applied to compare pairs of means and data are shown as the mean ± SEM of three independent experiments. *P < 0.05; ** P < 0.01; ***P < 0.001, arrow: the clump size of SSCs.

As the testicular EVs were secreted by spermatogonia that might interact physically with SSCs. Firstly, to assess the identity of the isolated SSCs, immunofluorescence revealed that more than 95% of isolated cells were positive for PLZF, OCT4, and MVH (supplementary Fig. S1), Reverse transcription-PCR and real-time PCR further showed that the isolated cells expressed the transcripts of PLZF, OCT4, MVH, Gfra1, and Etv5 (supplementary Fig. S2), collectively, these data suggest that the isolated cells are mouse SSCs phenotypically. to determine whether the testicular EVs could be taken up by SSCs, we labeled them with Exo-red, an acridine orange based dye that binds to RNA. One hour after exposure to testicular EVs, a remarkable uptake of the EVs by SSCs was observed, while the MEF feeder cells did not exhibit red fluorescence (Fig. 3D), and the supernatant of the labeled testicular EVs treated with Exo-red also did not stain the SSCs (**supplementary** Fig. S3). This indicated that SSCs might specifically take up testicular EVs by receptor-mediated endocytosis. To explore the effects of testicular EVs on the SSCs, the SSCs were exposed to testicular EVs from P8, P20, and P35, the result shown that the inhibitory effect of P20 EVs to the SSC proliferation was much better than P8 EVs (supplementary Fig.S4), so the testicular EVs from P20 were chose to further study. Next, the SSCs were exposed to various volumes of testicular EVs (0, 2.5, 5, and 10 µl), and CFA assays were performed (Yeh et al., 2007). As shown in Figure 3E, F, testicular extracellular vesicle suppressed SSC proliferation in a concentration-dependent manner and reduced the clump size of SSCs (Fig. 3E and supplementary table S1; arrow). These results suggest that testicular EVs can be taken up by SSCs and suppress their proliferation.

### 3.4 Thy1 positive EVs suppress the proliferation of SSCs cultured in vitro but not lead to apoptosis

To specifically separate the EVs from spermatogonia, we developed an approach designed to purify EVs bearing the undifferentiated A spermatogonia maker Thy1 (Figure4A), the use of magnetic beads directly conjugated to capture antibody, and the addition of the beads directly to testicular EVs samples. The Thy1 positive EVs were proved to be rich in Thy1, CD9 and Gfra1(another undifferentiated A spermatogonia maker) (Figure 4B, C). the effects of Thy1 positive and negative EVs on proliferation of SSCs were investigated by an EdU fluorescence assay. Notably, Thy1 positive EVs significantly repressed the proliferation of the SSCs, and the Thy1 negative EVs have no effect on the proliferation of the SSCs (Figure 4D, E). We next explored the Thy1 positive and negative EVs on apoptosis by an Annexin V-FITC/PI staining assay, the SSCs were exposure to Thy1 positive or negative EVs, the results showed that both Thy1 positive and negative EVs have no effect on the apoptosis in the SSCs (Figure 4F, G). In summary, Thy1 positive EVs suppress the proliferation of SSCs but not lead to apoptosis.

**Figure. 4.**
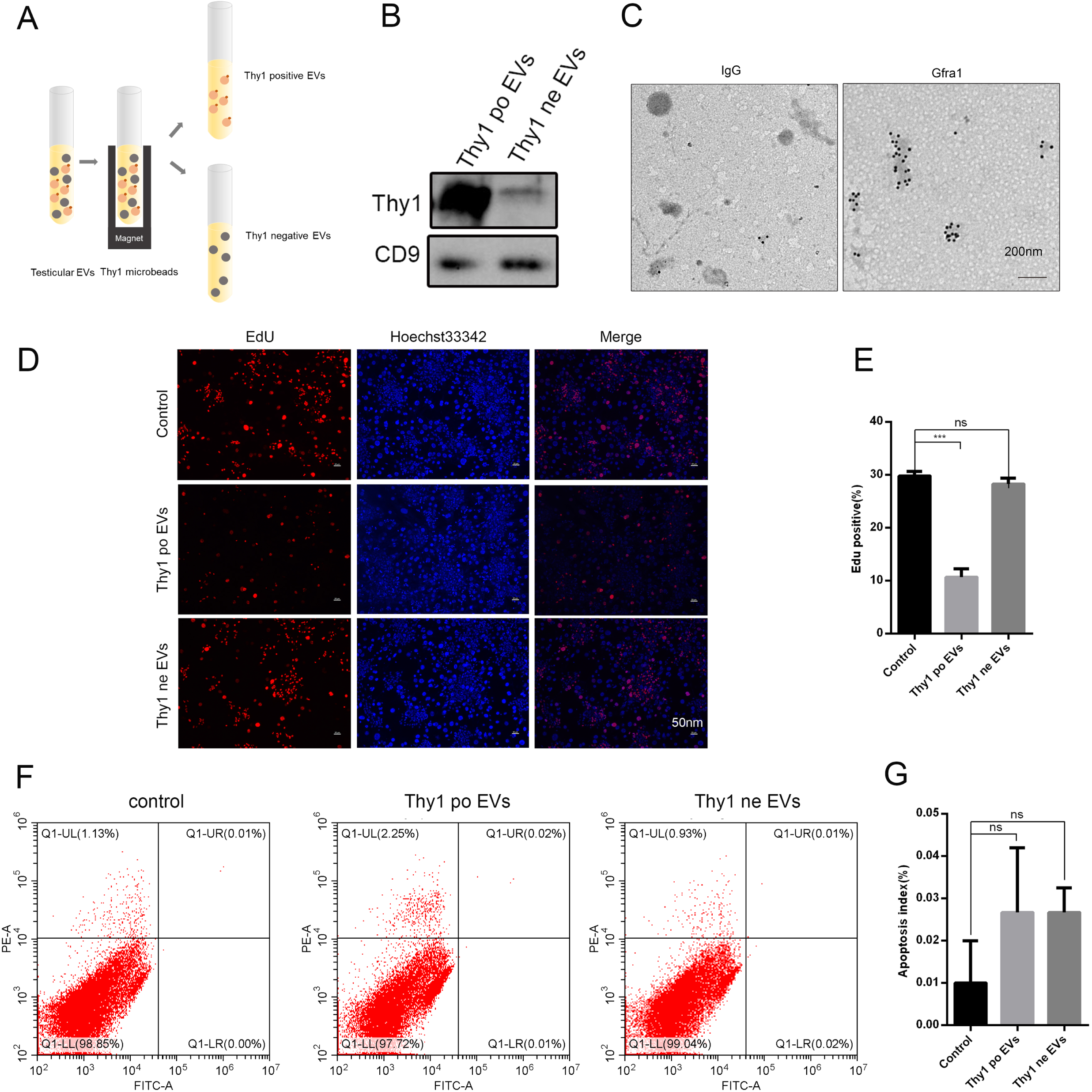
Testicular EVs from Thy1 spermatogonia suppress the proliferation of SSCs in vitro. (A) Schematic illustration of the direct immunoaffinity capture procedure Magnetic beads directly conjugated to anti-Thy1 were added directly to testicular EVs after ultracentrifugation. (B) Western blot analysis for Thy1 and CD9 using extracts from Thy1 positive EVs and Thy1 negative EVs. (C) Detection of Thy1 in Thy1 positive EVs by immunoelectron microscopy; IgG was used as a negative control. (D, E) Thy1 positive EVs regulates SSCs proliferation activity. Thy1 positive EVs enhanced BrdU incorporation, whereas Thy1 negative EVs did not affect cell proliferation activity. E, Histogram of data expressed as EdU positive cell index. Results are expressed as mean ± SEM from three independent experiments. ns > 0.05; *, P < 0.05; **, P < 0.01; ***, P < 0.005 (Student’s t test). (F, G) Evaluation of apoptosis in SSCs by Annexin V/PI assay (flow cytometry) treated with control Thy1 microbeads, Thy1 positive EVs and Thy1 negative EVs. F. Flow cytometry dot plots. G. Histogram of data expressed as apoptosis index. The bars represent means ± SD of three independent experiments. ns < 0.05.

## 4 DISCUSSION

EVs, are known as potent vehicles of intercellular communication by transferring proteins, lipids and nucleic acids both in prokaryotes and eukaryotes, thereby influencing various physiological and pathological process, for example, in cancer, the immune response angiogenesis and tissue regeneration (Merino-Gonzalez et al., 2016; Yanez-Mo et al., 2015). Whether EVs are involved in spermatogenesis remains uncharacterized. Here for the first time we have demonstrated one role of extracellular vesicle in spermatogenesis. First, TEM imaging, immunoelectron microscopy and western blot analysis revealed that spermatogonia secrete a large number of testicular EVs close to the basement membrane of seminiferous tubules: a feature that is common among mouse, rat, rabbit and human testes. Second, Exo-red labelling of testicular EVs and CFA assays showed that testicular EVs were specifically taken up by SSCs and repressed their proliferation *in vitro*. Finally, the testicular EVs were divided into the Thy1 positive EVs and Thy1 negative EVs, only the Thy1 positive EVs suppress the proliferation of SSCs. Thus, our study provides evidence that testicular EVs secreted by Thy1 spermatogonia play a significant role in regulating SSC proliferation

We separated the testicular EVs from testes at different stages of spermatogenesis by two-step enzyme digestion. To exclude EVs from interstitial cell, only the enzyme-digested supernatants of seminiferous tubules was used. The expressions of *GFRα1, Thy1, CD9*, and *c-Kit* in testicular EVs from the P8, P21, P35, 8Ws testes were detected by western blot (*CD81* was expressed in all testicular cells, as shown in Supplementary Fig. S5). In addition, we purified the EVs from type A spermatogonia by undifferentiated spermatogonia maker Thy1 (Figure4 A, B, C), suggesting the spermatogonia could secret the EVs in testis.

What is the significance of secreting EVs closed to basement membrane? Here we found that SSC proliferation was repressed when treated by testicular EVs or Thy1 positive EVs *in vitro*. This suggests that testicular EVs might play an important role in negatively regulating SSC proliferation (Fig. 3E, 3F, 4D, 4F), and which also could be related to the culture conditions. According to our TEM analysis, about 83% (40/48) testicular EVs interact physically with spermatogonia, whereas only 17% (8/48) interact with Sertoli cells, so whether the testicular EVs can be taken up by Sertoli cells and influence them needs further study.

## Supporting information

Figure S1. Immunocytochemical analysis of spermatogonial stem cell markers (OCT4, PLZF and MVH) was performed with mouse SSC clumps.

Figure S2. RT-QPCR analysis of spermatogonial stem cell makers (PLZF, OCT4, MVH, GFRA1, and ETV5) in the cultured mouse SSC clumps. GAPDH was used as

Figure S3. Fluorescence microscopy images of SSCs after treatment with the supernatant of the labeled testicular EVs.

Figure S5. Immunohistochemical analysis of CD81 in the mouse testes.

## ADDITIONAL INFORMATION

F.S., J.W., and Y.L. designed the study and wrote the paper. Y.L., and H.Y. performed the testicular extracellular vesicle-related experiments. F.Q., X.W.G and Y.J.W. performed the SSC-related experiments. Y.L., Y.J.W., A.J.L., Y.Z., and G.S.W. performed data analysis, F.Q, and Y.L. revised the manuscript according to reviewers’ suggestions.

## Supplemental data legends

Figure S2. RT-QPCR analysis of spermatogonial stem cell makers (PLZF, OCT4, MVH, GFRA1, and ETV5) in the cultured mouse SSC clumps. GAPDH was used as an experimental control.

Figure S4. The effect of testicular EVs from 8D, 20D and 35D on SSCs after 7 days were evaluated from cluster counts. Student’s t tests were applied to compare pairs of means and data are shown as the mean ± SEM of three independent experiments. *P < 0.05; ** P < 0.01; ***P < 0.001.

Figure S5. Immunohistochemical analysis of CD81 in the mouse testes.

