## Supplementary material for "Mouse thy1-positive spermatogonia suppress the proliferation of spermatogonial stem cells by Extracellular vesicles in vitro": Figure S1. Immunocytochemical analysis of spermatogonial stem cell markers (OCT4, PLZF and MVH) was performed with mouse SSC clumps.

**Supplementary Table S2. The cluster numbers of SSCs exposed to increasing concentrations of testicular EVs after 7 days**

| | 0 ul TEVs | 2.5ul TEVs | 5ul TEVs | 10ul TEVs |
|---|---|---|---|---|
| 1 | 141 | 96 | 83 | 71 |
| 2 | 111 | 110 | 71 | 73 |
| 3 | 123 | 98 | 96 | 71 |
| 4 | 112 | 102 | 73 | 57 |
| 5 | 112 | 93 | 79 | 60 |
| 6 | 117 | 113 | 71 | 71 |
