## Supplementary figures and images for "Mouse thy1-positive spermatogonia suppress the proliferation of spermatogonial stem cells by Extracellular vesicles in vitro"

### Figure S2. RT-QPCR analysis of spermatogonial stem cell makers (PLZF, OCT4, MVH, GFRA1, and ETV5) in the cultured mouse SSC clumps. GAPDH was used as

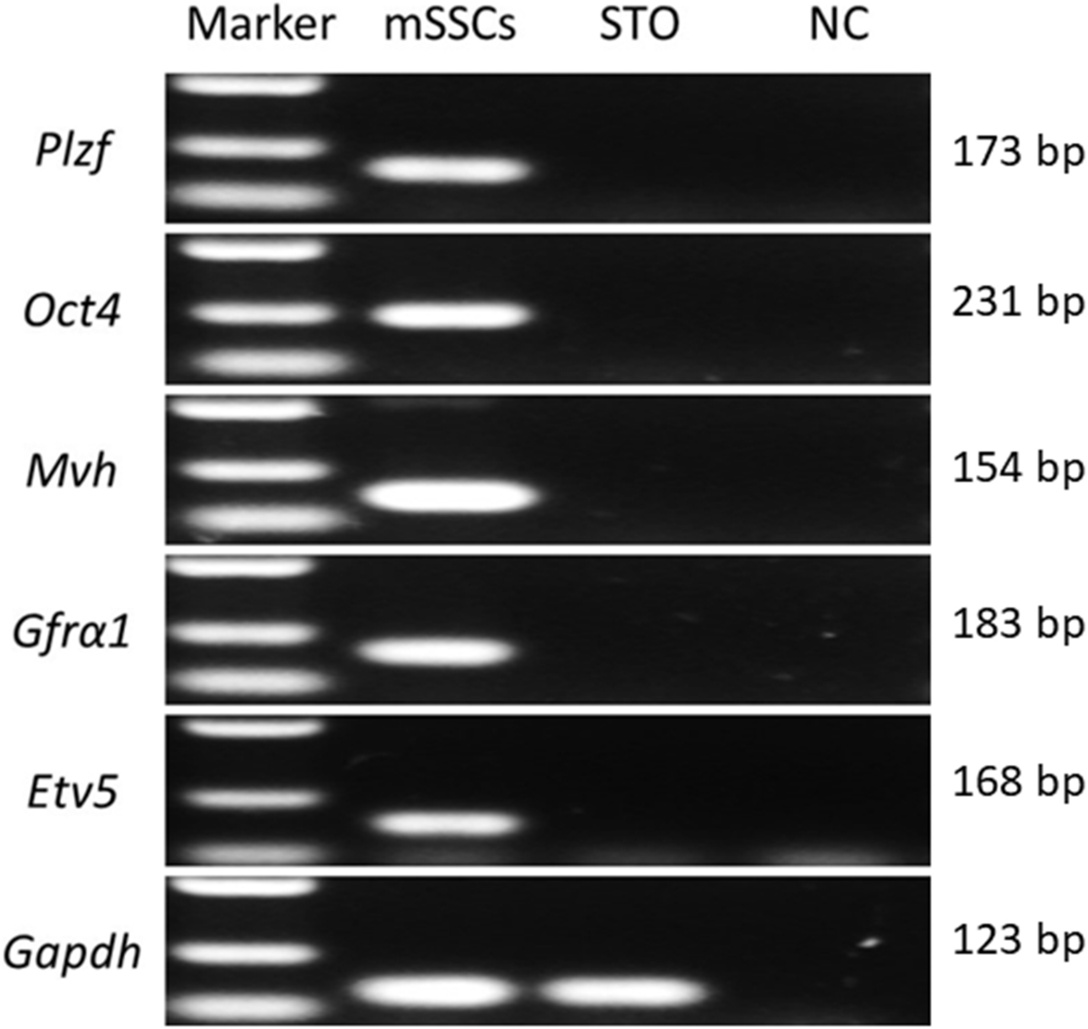


Note: NC represents negative control

### Figure S3. Fluorescence microscopy images of SSCs after treatment with the supernatant of the labeled testicular EVs.

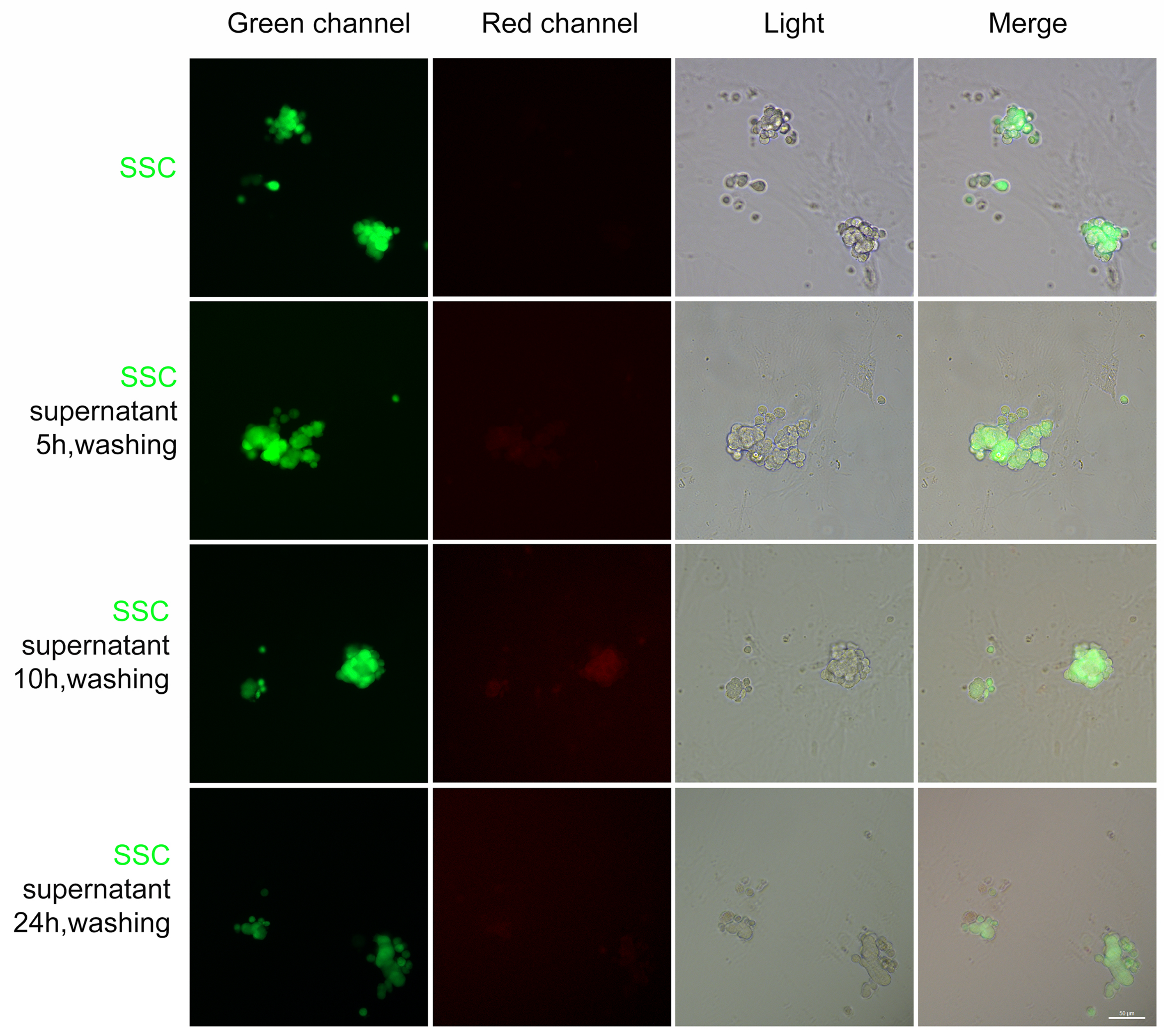

### Figure S5. Immunohistochemical analysis of CD81 in the mouse testes.

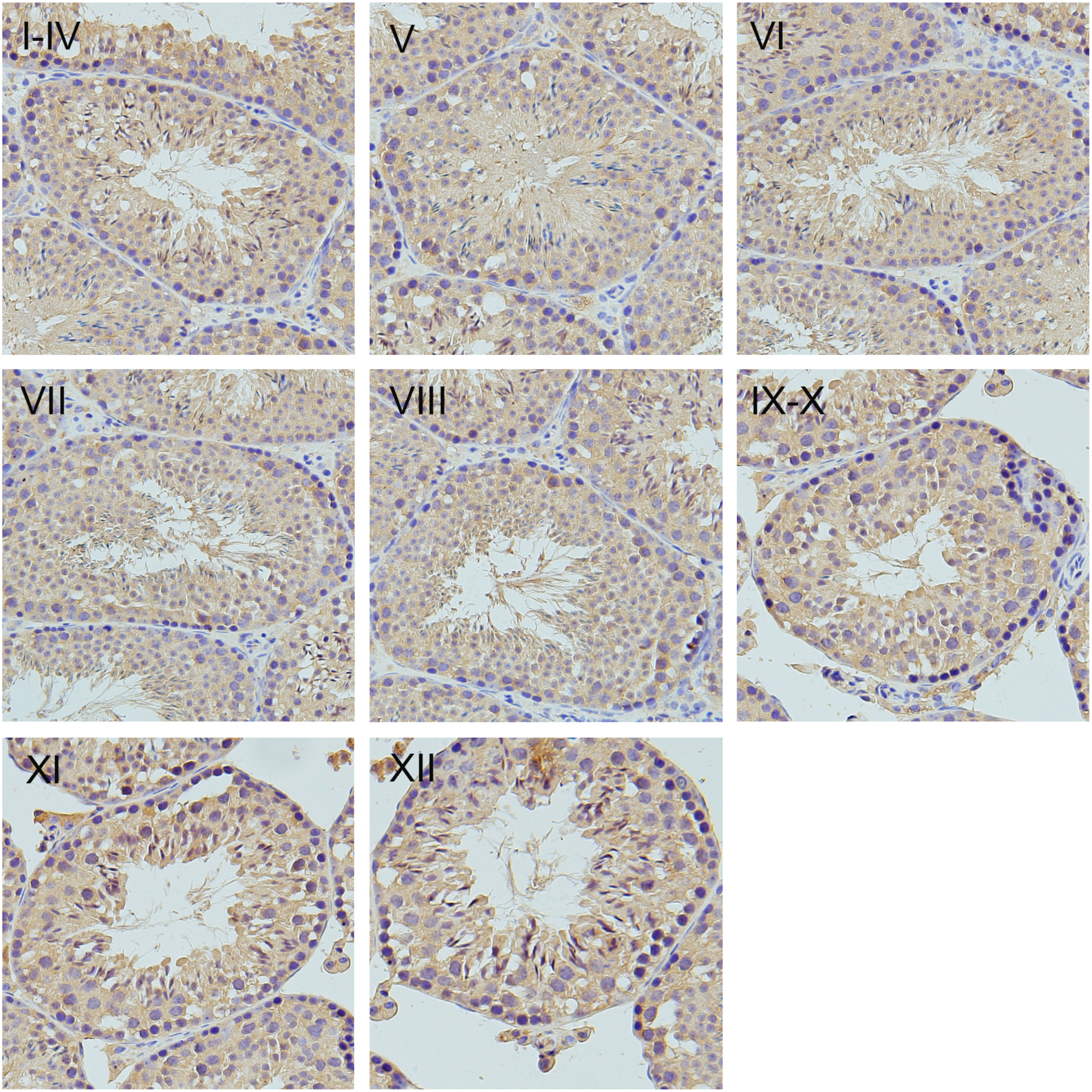

### Supplemental Data 1

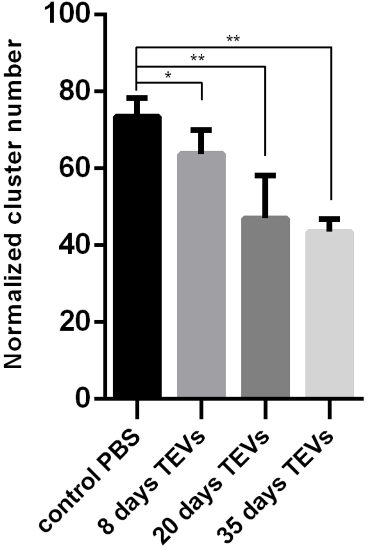
